# Dehydrojuncusol, a Natural Phenanthrene Compound Extracted from *Juncus maritimus* Is a New Inhibitor of Hepatitis C Virus Replication

**DOI:** 10.1101/469361

**Authors:** Marie-Emmanuelle Sahuc, Ramla Sahli, Céline Rivière, Véronique Pène, Muriel Lavie, Alexandre Vandeputte, Priscille Brodin, Arielle R. Rosenberg, Jean Dubuisson, Riadh Ksouri, Yves Rouillé, Sevser Sahpaz, Karin Séron

## Abstract

Recent emergence of direct acting antivirals (DAAs) targeting hepatitis C virus (HCV) proteins has considerably enhanced the success of antiviral therapy. However, the appearance of DAA resistant-associated variants is a cause of treatment failure, and the high cost of DAAs renders the therapy not accessible in countries with inadequate medical infrastructures. Therefore, search for new inhibitors and with lower cost of production should be pursued. In this context, crude extract of *Juncus maritimus* Lam. was shown to exhibit high antiviral activity against HCV in cell culture. Bio-guided fractionation allowed isolating and identifying the active compound, dehydrojuncusol. A time-of-addition assay showed that dehydrojuncusol significantly inhibited HCV infection when added after virus inoculation of HCV genotype 2a (EC_50_ = 1.35 µM). This antiviral activity was confirmed with a HCV subgenomic replicon and no effect on HCV pseudoparticle entry was observed. Antiviral activity of dehydrojuncusol was also demonstrated in primary human hepatocytes. No *in vitro* toxicity was observed at active concentrations. Dehydrojuncusol is also efficient on HCV genotype 3a and can be used in combination with sofosbuvir. Interestingly, dehydrojuncusol was able to inhibit replication of two frequent daclatasvir resistant mutants (L31M or Y93H in NS5A). Finally, resistant mutants to dehydrojuncusol were obtained and showed that HCV NS5A protein is the target of the molecule. In conclusion, dehydrojuncusol, a natural compound extracted from *J. maritimus*, inhibits infection of different HCV genotypes by targeting NS5A protein and is active against HCV resistant variants frequently found in patients with treatment failure.

**Importance:** Tens of millions of people are infected with hepatitis C virus (HCV) worldwide. Recently marketed direct acting antivirals (DAAs) targeting HCV proteins have enhanced the efficacy of the treatment. However, due to its high cost, this new therapy is not accessible to the vast majority of infected patients. Furthermore, treatment failures have also been reported due to appearance of viral resistance. Here we report on the identification of a new HCV inhibitor, dehydrojuncusol that targets HCV NS5A and is able to inhibit replication of replicons harboring resistance mutations to anti-NS5A DAAs used in current therapy. Dehydrojuncusol is a natural compound isolated from *Juncus maritimus*, a halophilic plant species very common in the coastlines worldwide. This molecule might serve as a lead for the development of new therapy more accessible to hepatitis C patients in the future.

## INTRODUCTION

Hepatitis C is a major cause of chronic hepatitis often associated with complications, such as liver cirrhosis and hepatocellular carcinoma (1, 2). A recent report estimates that approximately 71.1 million people are infected with hepatitis C virus (HCV) worldwide (3). To date no vaccine is available against HCV, mainly due to the diversity of viral isolates (4). In recent decades, various treatments have been established. The first direct-acting antiviral agents (DAA), targeting viral proteins were released in 2011. Since then, many DAA against NS3/4A, NS5A and NS5B have been marketed, with high rates of sustained viral response against all HCV genotypes and very few side effects (5). However, these treatments are very expensive and not accessible to all people infected (6). Moreover, there is a risk of selecting viral variants resistant to DAA leading to failure of the therapy, which will require new treatments (7).

HCV, a member of the *Flaviviridae* family, is an enveloped, single stranded RNA virus (8) encoding a single polyprotein which is cleaved co- and post-translationally. Non-structural proteins are involved in the replication of the viral genome and the production of new infectious particles in infected cells. The HCV life cycle can be divided into three major steps: entry, replication, and assembly/release. At each step, different sets of viral proteins and host factors are involved. The replication of the RNA genome takes place in the ‘membranous web’ which is composed of endoplasmic reticulum rearranged membranes (9). The replication complex includes viral proteins NS3/4A, NS4B, NS5A and NS5B.

The current hepatitis C therapy is very efficient but leads to the appearance of resistant associated substitutions (RASs). Some of the RASs are specific to a DAA, this is the case for NS5B RASs but some of them appear independently of the DAA used, especially for NS3/4A and NS5A RASs (10). Patients with NS5A resistant variant are very difficult to treat and retreatment with a combination of anti-NS5B and anti-NS5A antivirals does not always lead to a highly sustained viral response. Moreover, these mutated viruses are persistent for years in the blood serum (7). Lack of alternative therapy might be a problem for these patients in the future.

Natural products from plant species maintain a strong position in the drug discovery (11). A number of metabolites found in plants, among which are the phenolic compounds, are being cited as antimicrobials and resistance-modifying agents (12, 13). Natural products often remain a source of inspiration for medicinal chemistry, with semisynthetic modifications, or pharmacophore with natural origin (14). Numerous DAA from natural origin are now described and are able to target different steps of the HCV infectious cycle (15). Crude extracts from plants used in traditional medicine are also promising source of antiviral molecules. However, only silymarin, a standardized extract of Milk thistle (*Silybum marianum* (L.) Gaertn., Asteraceae) and some of its metabolites, silibinin A and B, a mixture of two diastereoisomers of flavonolignans, are developed in clinical trials to be used in hepatitis C infected patients (16). Extremophile plants either xerophytes (growing in arid climate), or halophytes (growing in saline soils) are natural reservoirs of rare bioactive compounds. By several molecular mechanisms, extremophile plants can resist to abiotic stresses and to infections by producing phytoalexins (17). Rare compounds with unique or common chemical structure present in extremophile plants might help to identify new DAA against HCV with unexpected mechanism of action.

It is estimated that the new DAA therapy against hepatitis C will benefit to a very small part of the patients (18). The use of plant extracts or compounds isolated from plant extracts should render the therapy more accessible to many patients in developing countries. Therefore, the search for such compounds and extracts is still needed to treat the vast majority of the patients in the coming decades. In this context, we have screened halophilic and xerophytic plant extracts from Tunisia for their anti-HCV activity. This led us to identify *Juncus maritimus* rhizome extract for its capacity to inhibit HCV infection (19). By use of a bio-guided fractionation approach, dehydrojuncusol, the active compound inhibiting HCV infection was identified and its mechanism of action against HCV replication characterized.

## RESULTS

### Dehydrojuncusol present in the methylene chloride partition of *J. maritimus* rhizome extract is an inhibitor of a post-entry step of HCV infection

We have previously screened sixteen plant extracts from eight different Tunisian extremophile plants for the presence of antiviral compounds (19). The strongest antiviral activity was observed for *Juncus maritimus* rhizome extract, a halophyte belonging to Juncaceae family. The methylene chloride partition of *J. maritimus* rhizome extract was the most active against HCV (19). To go further in the characterization of antiviral activity and identify active compound(s) a bioguided fractionation was performed leading to the isolation of two major phenanthrene derivatives (compounds 1 and 2). The chemical structures of these two major compounds were determined by comparison of their spectroscopic data (NMR and MS) with literature values. Compounds 1 and 2 were identified as two known phenanthrene derivatives, respectively juncusol and dehydrojuncusol (20) (Figure 1A). The purity of these compounds was checked by LC-UV-DAD (juncusol: 99.5%; dehydrojuncusol: 98.8%). Their antiviral activity was determined against HCVcc of genotype 2a. A time-addition assay was performed with the purified compounds at 10 µg/mL (corresponding to 37.5 and 37.8 µM for juncusol and dehydrojuncusol respectively) added either during virus inoculation, or post-inoculation, or continuously during both steps. Epigallocatechin gallate (EGCG), an inhibitor of HCV entry (21, 22) and Boceprevir, an inhibitor of NS3/4A protease, were used as controls. As shown in Figure 1B, juncusol and dehydrojuncusol were both active against HCV, dehydrojuncusol being the most active in the different conditions tested, co-addition, post-inoculation and continuously during infection. Only juncusol significantly decreased the number of cells (Figure 1C) showing limited cytotoxicity. Juncusol inhibited HCV infection at the post-inoculation step, whereas dehydrojuncusol was active both at inoculation and post-inoculation steps. Dehydrojuncusol was the most active at the post-inoculation step with more than 95% inhibition of HCV infection. As this compound was the most active, we decided to further analyze its antiviral activity.

**Figure 1.**
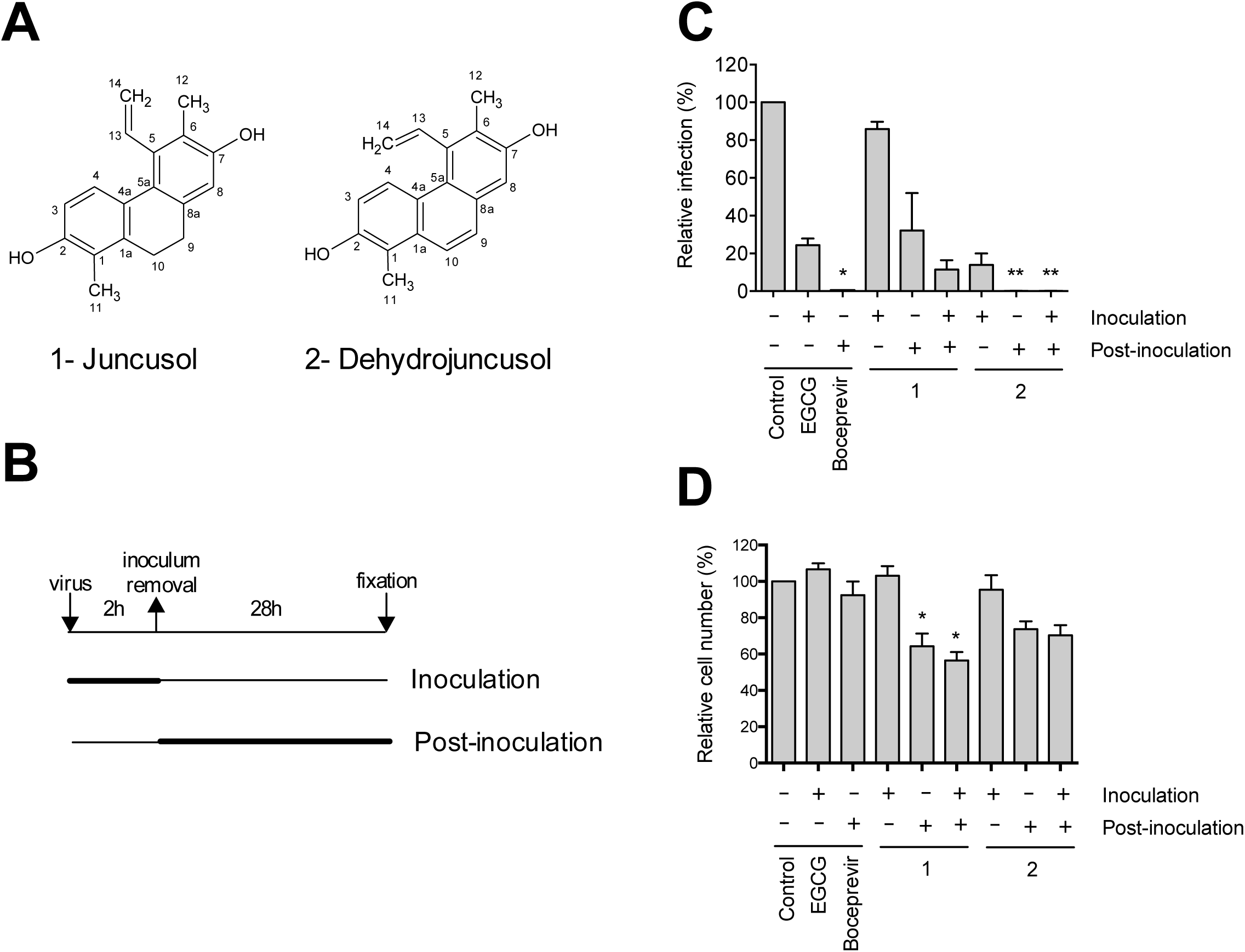
Anti-HCV activity of compounds isolated from *J. maritimus* rhizome extract. **(A)** Chemical structure of juncusol (**1**) and dehydrojuncusol (**2**). As represented in panel (B), infection was separated in different steps. Huh-7 cells were inoculated with HCVcc for 2h either in the presence (+) or not (-) of isolated compounds at 10 µg/mL (corresponding to 37.5 µM and 37.8 µM for juncusol and dehydrojuncusol, respectively), or 50 µM EGCG, or 1 µM boceprevir. DMSO 0.0001% was used as a control. Cells were further incubated in medium containing (+) or not (-) the compounds for 28h before fixation. **(C)** Infectivity was measured by the use of immunofluorescence labeling of HCV E1 envelope protein, and by calculating the number of infected cells. Data are expressed as a percentage of values measured with DMSO. **(D)** Nuclei were stained with DAPI to quantify the number of cells. Data are means of values obtained in 3 independent experiments performed in triplicate. Error bars represent standard error of the means (SEM). Significantly different from the control (DMSO): *, P<0.05; **, P<0.01.

The antiviral effect of dehydrojuncusol was examined by measuring HCV infectious titers. A dose-dependent decrease of the infectious titer was observed, with a 1-log_10_ decrease at 2.2 µM confirming its high anti-HCV activity (Figure 2A). A time-of-addition assay of dehydrojuncusol was performed and confirmed that this compound was significantly more active during a post-inoculation step (Figure 2B). No major effect of dehydrojuncusol was observed when cells were pre-incubated with the compound 2h before inoculation or when it was co-added with the virus. To measure the half maximal effective concentration EC_50_, a dose-response inhibition experiment was then performed with Huh-7 cells incubated with increasing concentrations of dehydrojuncusol at different steps of the infection, either during inoculation, or post-inoculation or continuously (Figure 2C). The results show that the EC_50_ of dehydrojuncusol was 1.35 µM when added continuously, 8.21 µM when added during inoculation, and 1.53 µM when added post-inoculation, confirming the major effect of the molecule at the post-inoculation step. The toxicity of the compound on Huh-7 cells was also tested in parallel at different time points, 24h, 48h, and 72h. The results showed that the cytotoxic concentration (CC_50_) of dehydrojuncusol was approximately 75.6 µM (Figure 2D), much higher than the active dose, yielding a selective index of 56.

**Figure 2.**
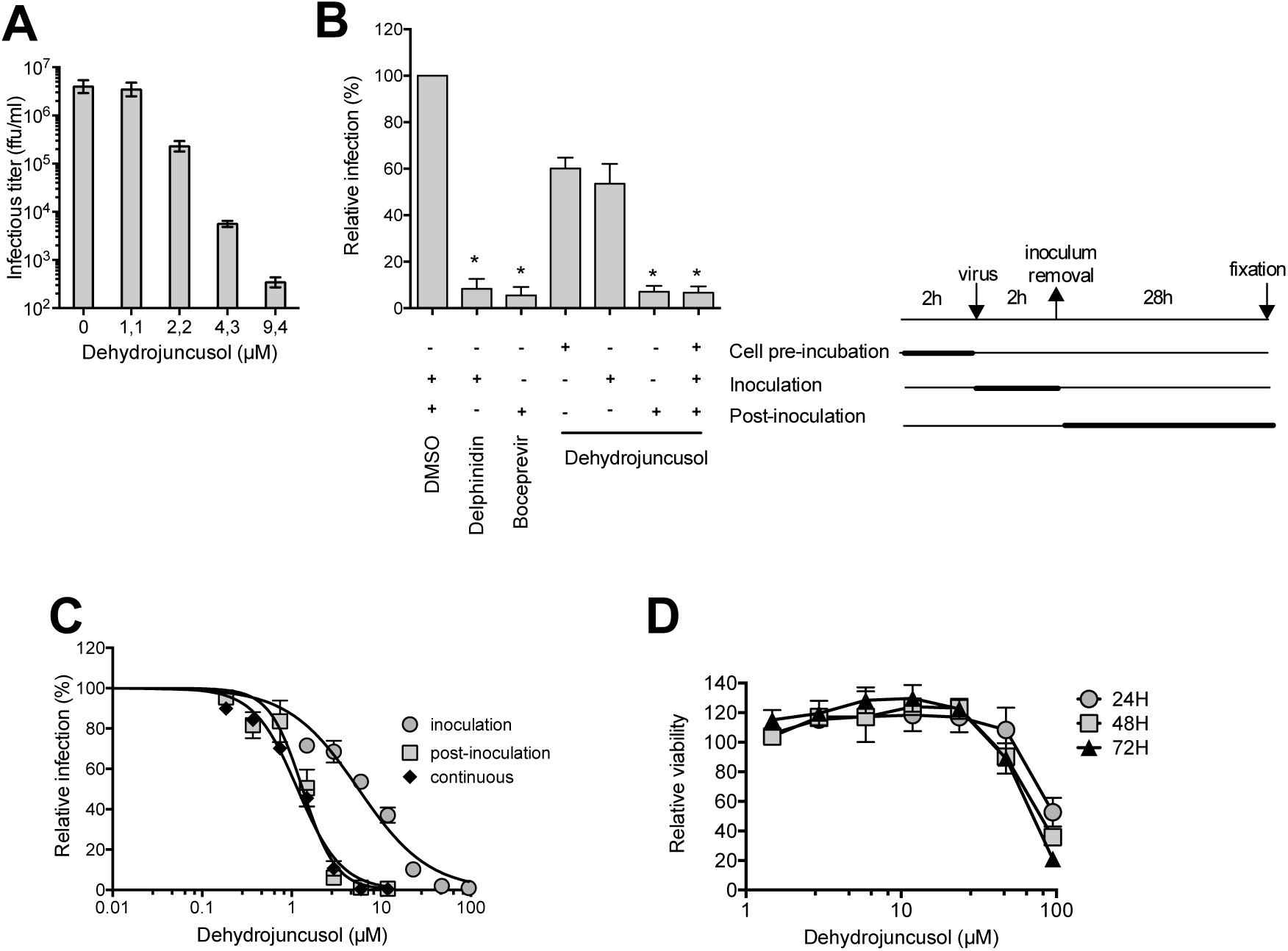
Dehydrojuncusol inhibits HCV infection in a dose dependent manner. **(A)** Huh-7 cells were infected with HCVcc in the presence of dehydrojuncusol at different concentrations. Supernatants were collected 48h post-inoculation and infectious titers (ffu/mL) were measured by serial dilutions of the supernatants and reinfection of naive Huh-7 cells for 48h. **(B)** Dehydrojuncusol at 3.8 µM, delphinidin at 50 µM or boceprevir at 1 µM were added at different time points as indicated (+) and represented on the right of the panel. Cells were either pre-incubated with the compounds for 2h, or compounds were added during HCVcc inoculation (2h) or post inoculation (28h). Infectivity was measured by the use of immunofluorescence labeling of HCV E1 envelope protein, and by calculating the number of infected cells. **(C)** Huh-7 cells were inoculated with HCVcc in the presence of increasing concentration of dehydrojuncusol either during the 2h inoculation, 28h post-inoculation or continuously. Infectivity was measured by the use of immunofluorescence labeling of HCV E1 envelope protein, and by calculating the number of infected cells. **(D)** Huh-7 cells were cultured in the presence of given concentrations of dehydrojuncusol. The viability was monitored using an MTS-based viability assay after 24h, 48h and 72h. Data are expressed as a ratio to control i.e. the condition without extracts. Data are means of values obtained in 3 independent experiments performed in triplicate. Error bars represent SEM. Significantly different from the control (DMSO): *, P<0.05.

Taken together, these results show that dehydrojuncusol inhibits predominantly a post-entry step of the HCV infectious cycle, most likely the replication, and, to a lesser extent, an early step of infection, potentially the entry step.

### Dehydrojuncusol inhibits replication of HCV but not entry

As shown above, dehydrojuncusol inhibits different steps of HCV infection. To determine if the molecule is able to inhibit HCV entry, HCV-pseudotyped retroviral particles (HCVpp) expressing E1E2 envelope glycoprotein of genotype 2a at their surface were produced and used to infect Huh-7 cells in the presence of dehydrojuncusol at different concentrations. Delphinidin at 50 µM, an inhibitor of HCV entry, was added as a control (23). A strong inhibition of HCVpp infection was observed with delphinidin. In contrast, no effect of dehydrojuncusol on HCVpp entry was observed even at concentration up to 94.7 µM (Figure 3A). This result shows that dehydrojuncusol is not an inhibitor of HCV entry.

**Figure 3.**
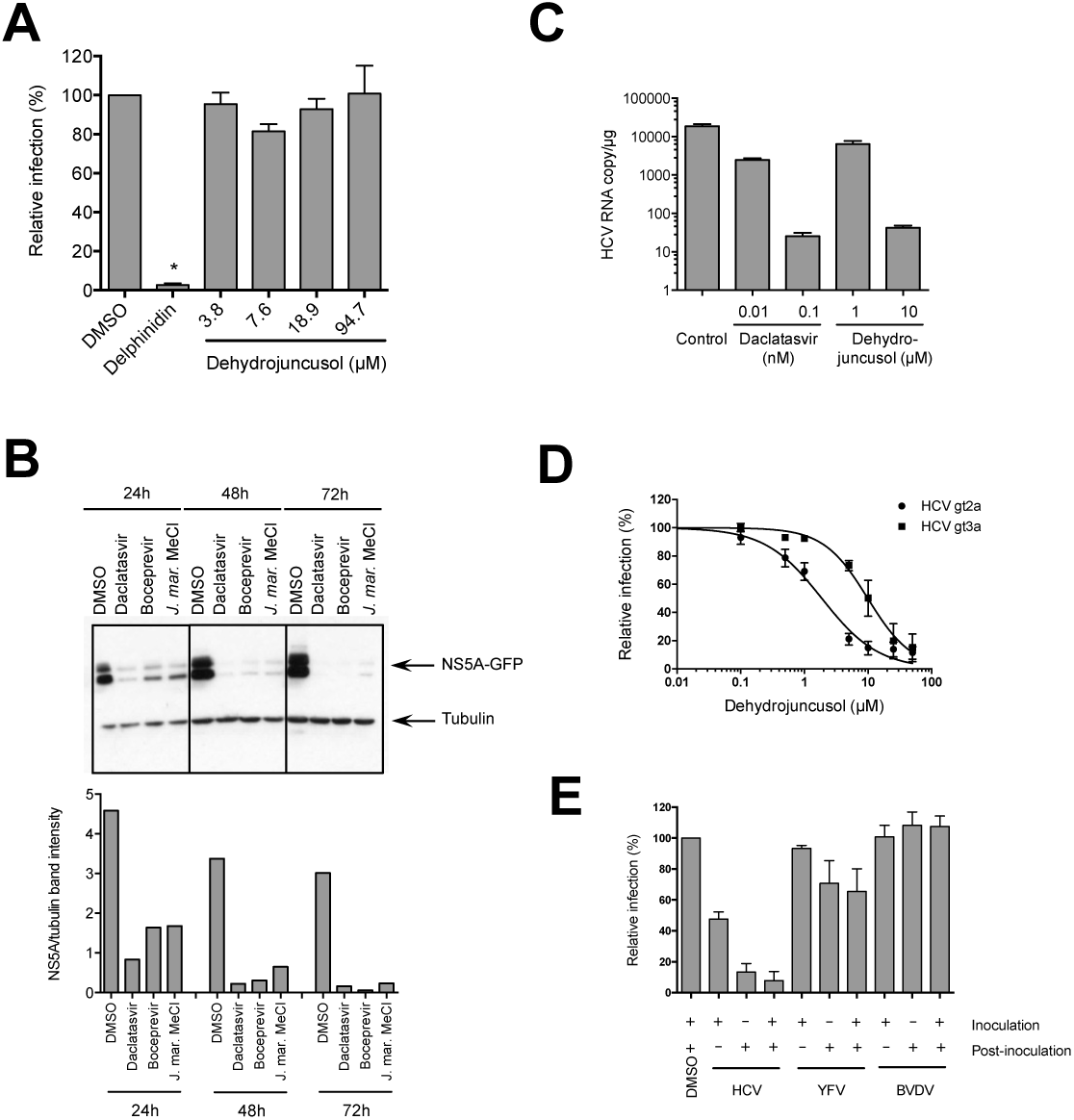
Dehydrojuncusol inhibits HCV replication. **(A)** HCVpp were used to inoculate Huh-7 cells in the presence of dehydrojuncusol at indicated concentrations. Delphinidin at 50 µM was used as a control. Cells were lysed 48h post-inoculation and luciferase activity quantified. Data are expressed as a ratio to the DMSO 0.0001 % control (100%). **(B)** Huh-7 cells stably expressing SGR-JFH1-NS5AGFP replicon were incubated for the indicated time with either 0.5 nM daclatasvir, 1 µM boceprevir, or 10 µg/mL of methylene chloride partition of *J. maritimus* (*J. mar.* MeCl). DMSO 0.0001 % was used as a control. Cells were lysed and lysates subjected to SDS-PAGE followed by western blot revelation of GFP and tubulin. The quantification of band intensity is shown in the graph **(C)** HCVcc-infected Huh-7 cells were maintained in the presence of dehydrojuncusol or daclatasvir at the indicated concentrations for 48h. Cells were lysed, RNA extracted, and HCV genomic RNA quantified by qRT-PCR. Data are representative of three experiments. Error bars represent SD. **(D)** Huh-7 cells were inoculated with HCVcc of genotype 2a (strain JFH1) and 3a (DBN3a) for 2h. Inoculum was removed and replaced with medium containing increasing concentration of dehydrojuncusol. Cells were fixed 30h post-infection and subjected to immunofluorescence staining with monoclonal anti-NS5A antibody (9E10). Data are means of values obtained in 3 independent experiments performed in triplicate. Error bars represent SEM. **(E)** Huh-7 cells were inoculated with HCVcc, or YFV and MDBK cells with BVDV in the presence (+) or absence (-) of 10 µg/mL of methylene chloride partition of *J. maritimus*. Inoculum was removed and replaced with medium with (+) or without (-) 10 µg/mL of methylene chloride partition of *J. maritimus*. Cells were fixed and infection detected by immunofluorescence labeling of viral proteins. Data are means of values obtained in 3 independent experiments performed in triplicate. Error bars represent SEM.

To test the activity of dehydrojuncusol on replication independently of entry, Huh-7 cells stably expressing a HCV replicon were used. This replicon, SGR-JFH1-NS5AGFP, expresses a GFP-tagged NS5A. Due to some difficulties to purify dehydrojuncusol in sufficient quantity from *J. maritimus* extract to perform all the tests, the methylene chloride partition containing dehydrojuncusol was used. The replicon cells were incubated with methylene chloride partition and lysed at different time points. Boceprevir and daclatasvir, two inhibitors of NS3/4A protease and NS5A protein, respectively, were used as controls. The activity of the molecules against HCV replication was determined by quantifying the expression of NS5A-GFP protein by western blots. As shown in Figure 3B, the expression of NS5A-GFP was strongly decreased in replicon cells incubated with boceprevir, daclatasvir and methylene chloride partition of *J. maritimus*, showing an inhibitory effect of dehydrojuncusol on HCV replication. Inhibition was observed at all time points, 24h, 48h and 72h of treatment, showing a rapid antiviral activity of methylene chloride partition. To confirm antiviral activity of dehydrojuncusol on the replication step, viral RNA quantification was performed, in HCV-infected Huh-7 cells treated for 48h with dehydrojuncusol or daclatasvir at different concentrations. A decrease of more than 2xLog_10_ of HCV genomic RNA level was observed with 10 µM dehydrojuncusol, similar to the decrease observed with daclatasvir, an NS5A inhibitor, at 0.1 nM (Figure 3C). Finally, to determine the effect of dehydrojuncusol on other genotypes, its antiviral activity was tested against a strain of genotype 3a. As shown in Figure 3D, dehydrojuncusol is able to inhibit HCVcc of genotype 3a with EC_50_ = 9.91 µM. Taken together, these results show that dehydrojuncusol is an inhibitor of HCV replication with an activity that is not limited to HCV genotype 2a.

### Dehydrojuncusol has no effect on bovine viral diarrhea virus (BVDV) and yellow fever virus (YFV) infection

To determine if the antiviral activity of dehydrojuncusol is specific to HCV, we tested the antiviral effect of the methylene chloride partition of *J. maritimus* against two other members of the *Flaviviridae* family, YFV and BVDV. As shown in Figure 3E, no significant decrease in infectivity was observed neither for YFV nor for BVDV after treatment with 10 µg/mL of *J. maritimus* methylene chloride partition. At this concentration, a strong inhibition of HCV infection was observed. This result shows that dehydrojuncusol is not active against all the members of the *Flaviviridae* family suggesting that it could target HCV protein(s) directly or a cellular factor specifically involved in HCV replication.

### Dehydrojuncusol inhibits HCV infection in PHH

Primary human hepatocytes (PHH) are a more relevant human pre-clinical model available to test the activity of an inhibitor on HCV infection. A dose-response experiment was performed in PHH inoculated with JFH-1 and treated with dehydrojuncusol for 28h after inoculation. The infectious titer of supernatants, negative-strand HCV RNA genome copy number and cytotoxicity of treatments were assayed 72h post-infection. As shown in Figure 4A, a dose-dependent decrease of HCV titer was observed with dehydrojuncusol showing that the molecule is able to inhibit HCV infection in PHH. Very interestingly, the EC_50_ of dehydrojuncusol in PHH was 1.14 µM, similar to the EC_50_ calculated in Huh-7 cells, 1.35 µM. The results were confirmed by the quantification of negative-strand HCV RNA genome (Figure 4B) that shows a similar decrease in the presence of dehydrojuncusol. The results presented in Figure 4C show that dehydrojuncusol is not cytotoxic in PHH at active concentrations.

**Figure 4.**
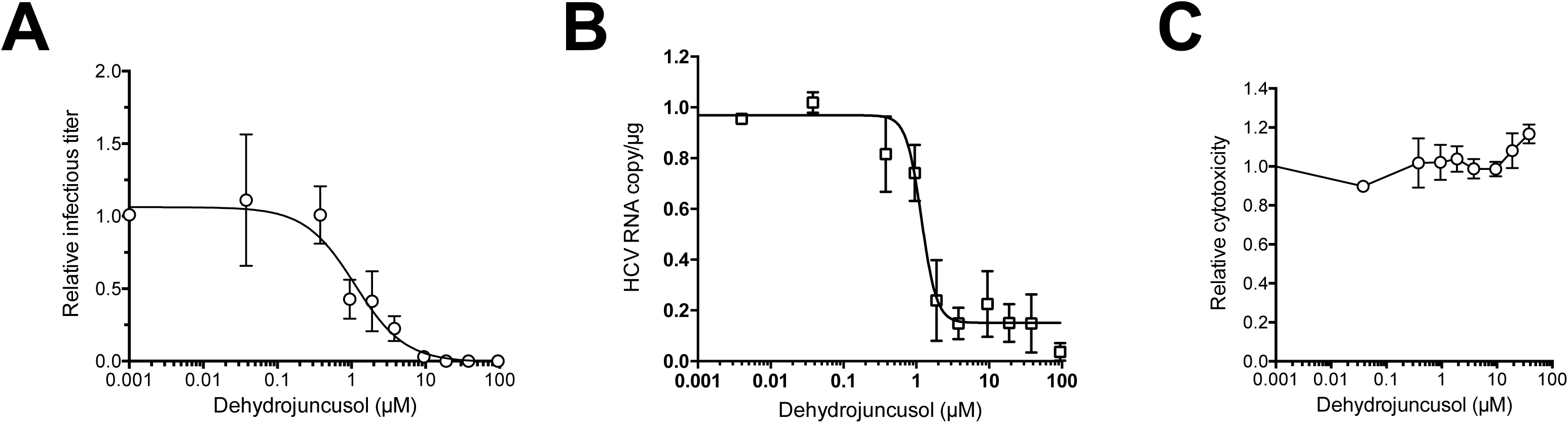
Dehydrojuncusol inhibits HCV infection in PHH. PHH were inoculated with HCVcc and incubated for 28h after inoculation in the presence of the indicated concentrations of dehydrojuncusol. Supernatants were harvested 72h post-inoculation for determination of infectivity (A) and cytotoxicity (C). Cells were lysed, RNA extracted and negative-strand HCV RNA genome copy number was quantified (B). Data are expressed as a ratio to carrier DMSO 0.0001 % control. Data are means (± SEM) of 3 independent experiments performed in triplicate.

### Dehydrojuncusol inhibits replication of HCV mutants resistant to NS5A inhibitors

Some patients treated with the newly approved DAA molecules targeting NS5A or NS5B become resistant to the therapy due to the appearance of HCV resistant mutants. Once present in patients, these resistant mutants are very difficult to eliminate, particularly for the resistant mutants selected during a treatment with anti-NS5A molecules, like daclatasvir, ledipasvir or ombitasvir. To determine if dehydrojuncusol could be active against HCV resistant mutants, we generated mutated replicons harboring point mutation L31M or Y93H in NS5A. These two mutations conferred resistance to NS5A targeting drugs, including daclatasvir, ledipasvir and ombitasvir, in patients infected with HCV of different genotypes, and are often associated with treatment failure (7, 10). Huh-7 cells harboring mutant replicons were incubated with methylene chloride partition of *J. maritimus*, containing dehydrojuncusol, and lysed after 72h of treatment. Daclatasvir and boceprevir, were used as controls. As shown in Figure 5A, in the L31M and Y93H replicon cell lysates, the NS5A-GFP protein was detected after 72h of treatment with daclatasvir showing that these replicons are resistant to daclatasvir at 0.5 nM. When treated with boceprevir, NS5A-GFP was not detected showing that the mutants are still sensitive to boceprevir, as expected. In cells expressing mutated replicons treated with methylene chloride partition at 10 µg/mL, NS5A-GFP was not detected with the L31M mutant, showing that dehydrojuncusol is able to inhibit the replication of a daclatasvir-resistant mutant. However, it was still detectable at lower level with the Y93H daclatasvir-resistant mutant, showing that this mutant is less sensitive to dehydrojuncusol than L31M. To quantify this antiviral activity, a dose-response experiment was performed on replicon cells treated with either daclatasvir or dehydrojuncusol and the number of cells expressing NS5A-GFP after 72h of treatment was quantified. As shown in Figure 5B, the L31M mutant was very sensitive to the dehydrojuncusol with an EC_50_ of 0.22 µM similar to wild-type 0.40 µM, whereas the Y93H daclatasvir-resistant was less sensitive to dehydrojuncusol than the wild-type replicon, (EC_50_ of 3.62 µM), correlating with the results obtained by Western blot. However, the EC_50_ fold change between wild-type and Y93H replicon was only of 9, much lower than the EC_50_ fold change observed with daclastasvir (Figure 5C), which was calculated to be over 100 for both mutants (24).

**Figure 5.**
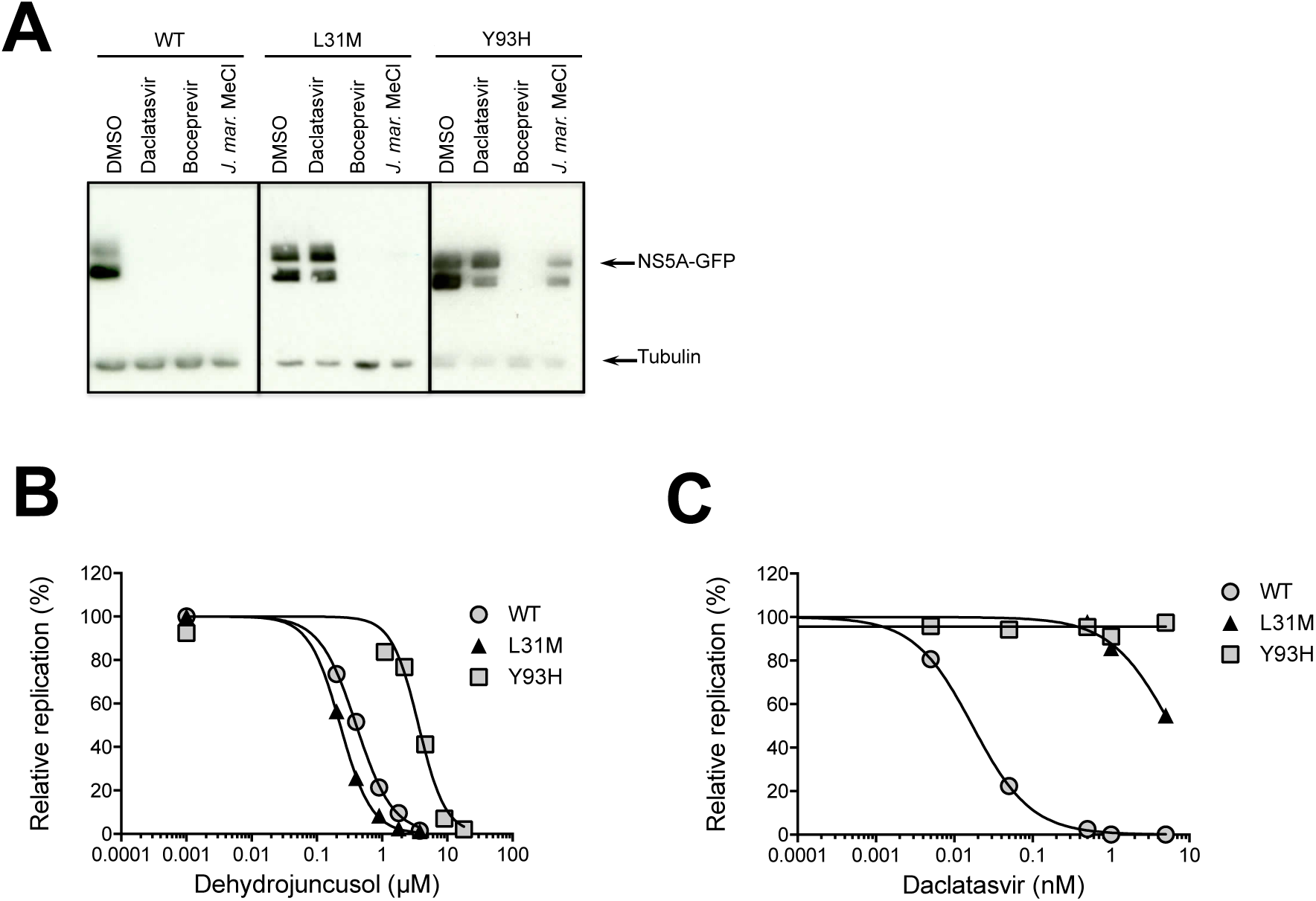
Dehydrojuncusol inhibits HCV replication of daclatasvir NS5A resistant mutants. **(A)** Huh-7 cells stably expressing subgenomic replicon SGR-JFH1-NS5AGFP wild-type or harboring L31M or Y93H mutation in NS5A that confer resistance to daclatasvir were incubated for 72h either with 0.5 nM daclatasvir, 1 µM boceprevir, or 10 µg/mL of methylene chloride partition of *J. maritimus* (*J. mar.* MeCl). DMSO 0.0001 % was used as a control. Cells were lysed and lysates subjected to SDS-PAGE followed by western blot revelation of GFP and tubulin. **(B)(C)** Huh-7 cells stably expressing subgenomic replicon SGR-JFH1-NS5AGFP wild-type or harboring L31M or Y93H mutation in NS5A that confer resistance to daclatasvir were incubated for 72h with increasing concentrations of dehydrojuncusol **(B)** or daclatasvir **(C)**. Cells were fixed and the percentage of GFP positive cells was quantified, as a measure of replication. Replication of subgenomic replicon was expressed as a ratio to carrier control DMSO 0.0001 %. Data are means of values obtained in 3 independent experiments performed in triplicate. Error bars represent SEM.

### The target of dehydrojuncusol is NS5A

In order to identify the viral target of dehydrojuncusol, JFH1 resistant mutants were generated by infecting Huh-7 cells with JFH1, followed by treatment with 5 × EC_50_ of dehydrojuncusol, for 3 weeks. Viral RNA was extracted every 3-4 days to follow the appearance of resistant viruses (data not shown). Viral RNA extracted at day 21 was sequenced and three mutations at the same amino-acid position of NS5A were detected, F28I, F28L, and F28V. These mutations were individually inserted in the JFH1 genome by reverse genetic. The three mutants exhibited a strong resistance to dehydrojuncusol, showing that the amino acid residue F28 is required for the anti-HCV activity of dehydrojuncusol (Figure 6A). In parallel, we also tested the resistance of these mutants toward daclatasvir. The F28I, F28L and F28V remained sensitive to daclatasvir (Figure 6B).

**Figure 6.**
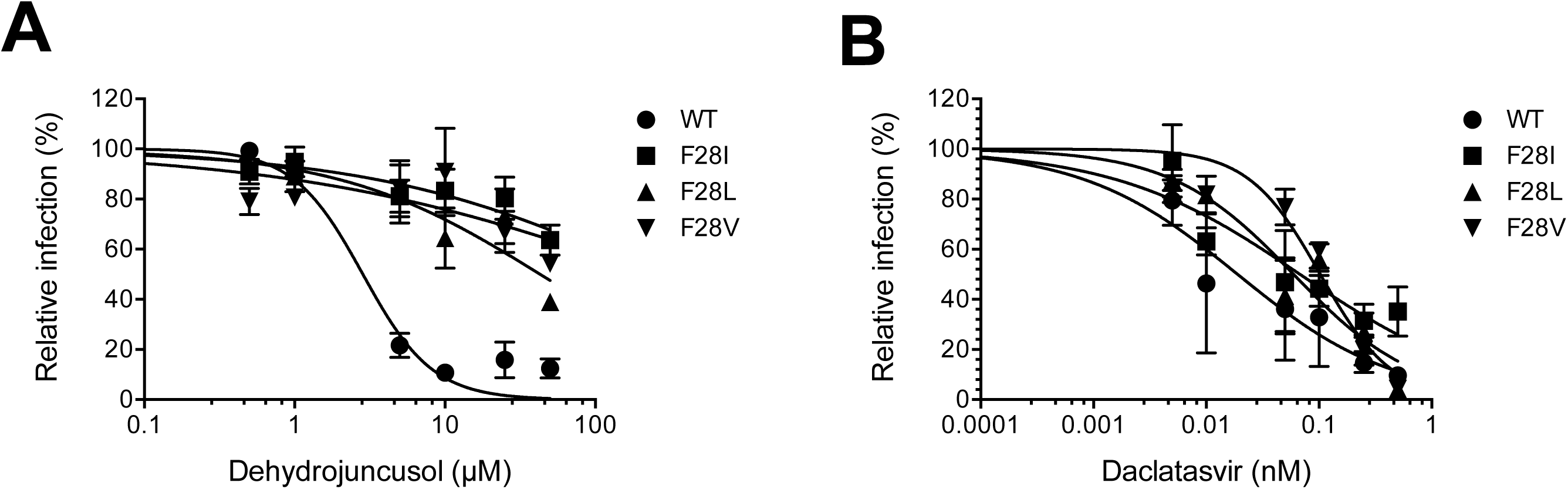
Dehydrojuncusol targets NS5A. Huh-7 cells were inoculated with wild-type HCVcc or HCVcc harboring F28I, F28L or F28V mutations in the presence of given concentrations of dehydrojuncusol (A) or daclatasvir (B). Cells were fixed 30h post-inoculation and infectivity was measured by the use of immunofluorescence labeling of HCV E1 envelope protein, and by calculating the number of infected cells. Data are expressed as a percentage of values measured with 0.0001 % DMSO. Data are means of values obtained in 3 independent experiments performed in triplicate. Error bars represent SEM.

### Dehydrojuncusol can be used in combination with Sofosbuvir

The results presented ahead show that dehydrojuncusol encompasses many characteristics of a DAA to be used in therapy. In order to determine the potential use of dehydrojuncusol in combination with the known DAA, combination study of dehydrojuncusol and sofosbuvir, an inhibitor of HCV polymerase NS5B (25), was performed. Dehydrojuncusol was added at different concentrations along with sofosbuvir at fixed concentrations, 400 and 600 nM, during infection of Huh-7 cells with HCVcc. The result shows that Sofosbuvir can increase the antiviral activity of dehydrojuncusol in an additive manner (Figure 7). The EC_50_ of dehydrojuncusol is decreased down to 1.10 nM in the presence of sofosbuvir at 600 nM, demonstrating that dehydrojuncusol could be used in combination with DAA used in hepatitis C therapy.

**Figure 7.**
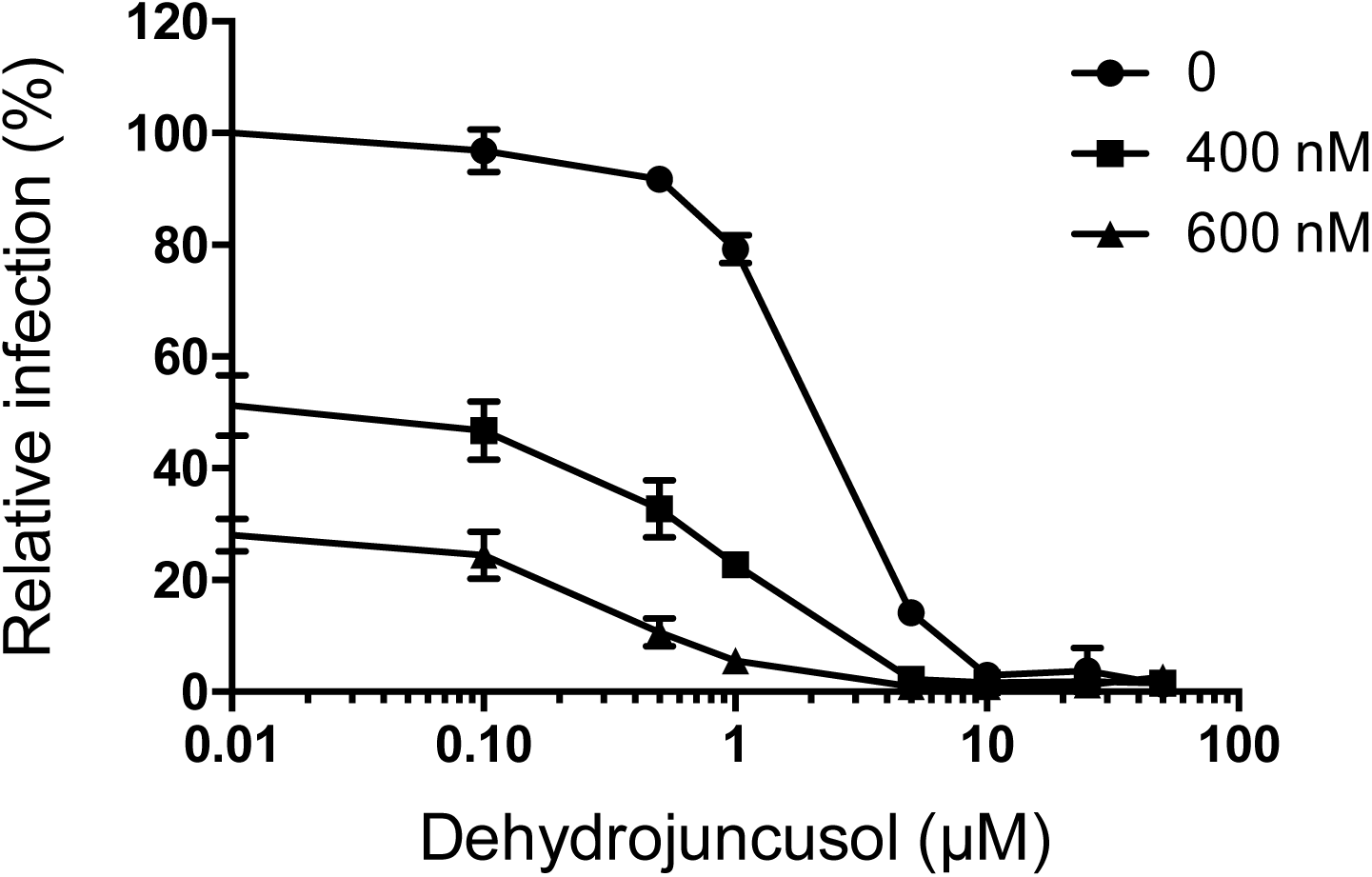
Dehydrojuncusol can be used in combination with sofosbuvir. Huh-7 cells were inoculated with HCVcc. Inoculum was removed and replaced by medium containing dehydrojuncusol at given concentrations combined with sofosbuvir at 400 or 600 nM. Cells were fixed 30h post-inoculation and infectivity was measured by the use of immunofluorescence labeling of HCV E1 envelope protein, and by calculating the number of infected cells. Data are means of triplicate values obtained in one experiment representative of 3 independent experiments. Error bars represent SD.

## DISCUSSION

Plants are known to be an important source of numerous bioactive compounds (14). Here, we highlighted the strong inhibition of HCV infection obtained in HCV in cell culture (HCVcc) system by a bioactive molecule, dehydrojuncusol, isolated from a *J. maritimus* rhizomes extract. More interestingly, we also showed that this natural product strongly inhibits the virus in PHH, which is one of the most relevant human pre-clinical model for HCV infection. Many different drugs are now available to cure hepatitis C, but in some patients, appearance of viral variants resistant to the treatment makes them difficult to cure, especially for NS5A resistant mutants. Unfortunately, the sets of marketed molecules available are not efficient to cure such resistant viruses (7). In this work, we demonstrate that dehydrojuncusol has the capacity to inhibit two NS5A resistant mutants appearing in patients treated with anti-NS5A inhibitors, L31M and Y93H, which are two major resistant variants appearing after treatment with daclatasvir, ledipasvir or ombitasvir (7, 10). These mutants are often associated with treatment failure. Therefore, dehydrojuncusol or molecules derived from dehydrojuncusol might be of great interest in the future to cure patients with resistant variants and treatment failure. Here, we demonstrate that dehydrojuncusol anti-HCV activity is not genotype specific since this molecule is able to inhibit HCV genotype 3a in addition to genotype 2a. Furthermore, we also show that dehydrojuncusol can be used in combination with sofosbuvir, the most widely used DAA that targets NS5B polymerase.

Even if dehydrojuncusol is able to inhibit replication of HCV replicon harboring point mutations in NS5A (L31M and Y93H), it seems likely that the viral target of this molecule is NS5A. Dehydrojuncsuol resistant HCV mutants were obtained and we demonstrated that a single mutation of NS5A F28 is responsible for the resistance to dehydrojuncusol. This amino acid residue was mutated into three different amino acids, isoleucine, leucine, and valine and each of them confers resistance to dehydrojuncusol. Phenylalanine 28 is one of the first amino-acid residues of NS5A domain 1. This domain is the target of daclatasvir, ombitasvir and ledispavir (26). Resistance mutations in NS5A at the position 28 have been already described for genotype 1 (M28) which confer resistance to daclatasvir, ombitasvir or ledipasvir (7, 10, 27, 28). For JFH1 genotype 2a, F28S resistance mutation has been observed for daclatasvir (24) and ombitasvir (29). In this study, no other mutation than the one at position F28 has been identified suggesting that dehydrojuncusol might interact with NS5A at a site different than daclatasvir or other known anti-NS5A inhibitors.

Unexpectedly for a replication inhibitor, dehydrojuncusol also inhibits HCV infection when it is added to the cells during virus contact and removed later on (Figure 2B and 2C). This ability to inhibit an early step of HCVcc infection is probably not due to an inhibition of HCV entry as this molecule is not active on HCVpp (Figure 3A). It is likely that this effect results from dehydrojuncusol molecules that enter cells during viral inoculation, remain trapped inside cells and are able to inhibit HCV replication afterwards.

The EC_50_ of dehydrojuncusol is close to 1 µM, in the replicon, in the HCVcc system, and in PHH, a concentration relatively low for a natural product. This molecule might serve as a starting point for a study of structure activity relationships and drug design to increase its antiviral capacity and reduce its EC_50_. It is rare that a compound is as active in Huh-7 as in PHH. The EC_50_ of dehydrojuncusol could even be decreased by 3xLog_10_ when it was combined with sofosbuvir at 600 nM. Toxicity of dehydrojuncusol in Huh-7 cells is relatively low with a CC_50_ of approximately 75 µM, leading to a selective index of 56. Similarly, no toxicity was observed in PHH even at high concentration.

Unfortunately, little is known about the bioavailability of this compound in animal models or in humans. More generally, few/no pharmacokinetics studies of phenanthrene derivatives *invivo* or in humans were reported. Natural phenanthrenes are an uncommon class of aromatic metabolites, which in a biosynthetic point of view can originate from stilbene precursors and more rarely from diterpenoid precursors. These compounds are biosynthesized by a limited number of botanical families. Orchidaceae and Juncaceae are the main botanical families known to produce this kind of natural products (30, 31). Phenanthrenes and 9,10-dihydrophenanthrenes showed a large panel of biological activities, including among others cytotoxicity, antimicrobial, spasmolytic and anti-inflammatory activities. However, this potential is not sufficiently investigated and not really exploited (31). Concerning the antiviral activities of phenanthrene derivatives, a limited number of studies exist in the literature. Some phenanthrene derivatives isolated from *Tamus communis* (Dioscoreaceae) showed antiviral activity against two RNA viruses: vesicular stomatitis virus (VSV) and human rhinovirus serotype 1B (HRV 1B) (32). So far, no antiviral activity was highlighted for dehydrojuncusol. Only a moderate anti-inflammatory activity has been shown for this natural product (33).

*J. maritimus* is a perennial plant which thrives in particular on saline soils. In the framework of our study, this halophyte was collected in Tunisia, but *J. maritimus* is present worldwide in the coast line of many different areas, including several European countries such as France. Few traditional uses are mentioned for this plant in Tunisia, with the exception of its use in making baskets and curtains. Its usage as an antiviral agent is a new opportunity for economic valorization of this plant. One of the disadvantages of our study is that the part of the plant used is not renewable (rhizomes), so the chemical synthesis of dehydrojuncusol or a more active derivative could be an alternative. It would be also interesting to search for dehydrojuncusol in other *Juncus* species and especially in parts of plant renewable, such as aerial parts. Interestingly, this phenanthrene derivative has been previously isolated from the aerial parts of *Juncus* species, such as *J*. *acutus* (34) and *J. effusus* (35), increasing its potential for its use in low income countries.

In conclusion, we showed that dehydrojuncusol, an active compound present in *J. maritimus* rhizomes, is a new DAA that targets HCV NS5A and inhibits replication of resistant viruses appearing in patients treated with anti-NS5A inhibitors and responsible for treatment failure. Plants should be more considered, in the future, as a source of antiviral agents and dehydrojuncusol could constitute an alternative treatment for patients resistant to currently available DAA.

## MATERIALS AND METHODS

### Chemicals

Dulbecco’s modified Eagle’s medium (DMEM), Opti-MEM, phosphate buffered saline (PBS), glutamax-I, fetal bovine serum were purchased from Invitrogen (Carlsbad, CA). 4’,6-Diamidino-2-phenylindole (DAPI) was from Molecular Probes (Thermo Fischer Scientific, Waltham, USA). EGCG was from Calbiochem (MerckChemicals, Darmstadt, Germany) and was >95% pure. Delphinidin chloride was from Extrasynthèse (Lyon, France) and was >96% pure. Daclatasvir (BMS-790052) and sofosbuvir (PSI-7977, GS-7977) were from Selleckchem (France). Stocks of EGCG and delphinidin were resuspended in dimethylsulfoxide (DMSO) at 0.5 M. Plant extracts were resuspended in DMSO at 25 mg/mL. Boceprevir was kindly provided by Philippe Halfon (Hôpital Européen, Laboratoire Alphabio, Marseille, France). Other chemicals were from Sigma (St. Louis, MO).

### Antibodies

The anti-E1 monoclonal antibody (A4) (36), anti-YFV envelope protein 2D12 antibody (ATCC CRL-1689) and anti-BVDV NS3 protein OSC-23 antibody (37) were produced *in vitro* by using MiniPerm apparatus (Heraeus, Hanau, Germany). The anti-NS5A, 2F6/G11 and 9E10 were from Austral Biologicals (San Ramon, USA) or kindly provided by C.M. Rice (Rockefeller University, NY), respectively. The mouse monoclonal anti-GFP antibody was from Roche. Mouse anti-β tubulin monoclonal antibody (TUB 2.1) was from Sigma. Cy3-conjugated goat anti-mouse IgG and peroxidase-conjugated goat anti-mouse IgG antibodies were from Jackson Immunoresearch (West Grove, PA, USA).

### Cells and culture conditions

Huh-7 and HEK 293T (ATCC number CRL-11268) were grown in DMEM supplemented with glutaMAX-I and 10% fetal bovine serum (complete culture medium), and Madin-Darby Bovine Kidney (MDBK; ATCC number CCL-22) in DMEM supplemented with glutaMAX-I and 10% horse serum in an incubator at 37°C with 5% CO_2_. The primary human hepatocytes were from Biopredic International (Saint-Grégoire, France) and maintained in primary culture as described previously (23, 38).

### Collection of plant, extraction and purification of natural products

*Juncus maritimus* rhizomes were collected in October 2013 from a coastal region in the North-East of Tunisia (Soliman) and a voucher specimen (184) was deposited at the Herbarium of the Laboratory of Extremophile Plants at the Biotechnology Centre (Technopark of Borj-Cédria). Dried and powdered rhizomes were soaked in methanol (15 mL/g; 1 × 24h; 2 × 48h) to afford 300.0 g of a crude methanolic extract. A part of the extract (215.0 g) was dissolved in water and then partitioned with methylene chloride (CH_2_Cl_2_) to afford a CH_2_Cl_2_ partition (15.0 g). Preparative Centrifugal Partition Chromatography (CPC) (Armen Instrument) was carried out on CH_2_Cl_2_ partition (1.45 g) using a quaternary biphasic solvent system Arizona X (n-Hept/EtOAc/MeOH/H_2_O; 9:1:9:1; v/v/v/v) in ascending mode for 30 min. Fifteen fractions (MCX1 to MCX15) were pooled according to TLC and UV analysis. Compounds 1 (22.2 mg) and 2 (25.9 mg) were isolated from MCX15 (200 mg) by preparative High Performance Liquid Chromatography (HPLC) (Shimadzu instrument), using the following elution program (EP1): 10-38% B (0-2 min), 38-45% B (2-8 min), 45-51% B (8-25 min), 51-96% B (25-26 min), 96%-96.1% B (26-29 min), 100% B (29-35 min).

### HCVcc

We used a modified JFH1 strain (Japanese Fulminant Hepatitis-1, genotype 2a) containing cell culture adaptive mutations (39, 40), and the HCV strain DBN3a of genotype 3a (41). JFH1 and DBN3a were kindly provided by T Wakita (National Institute of Infectious Diseases, Tokyo, Japan) and Jens Bukh (University of Copenhagen, Denmark), respectively. The viral stocks were produced in Huh-7 cells. Huh-7 cells were infected with a pre-stock of HCVcc in flasks. After 24h, 48h and 72h the supernatants of flasks were collected. The titer of the stock was 5.10^6^ and 5.10^5^ focus forming unit (ffu)/mL for JFH1 and DBN3a respectively. For infection assay, Huh-7 cells, 6,000/well, seeded in 96-well plate were inoculated with HCVcc at a multiplicity of infection (MOI) of 0.8 during 2h at 37°C then the inoculum was removed and cells were incubated in complete culture medium for 28h at 37°C. Compounds were added to cells either for 2h at 37°C before infection (preincubation condition), for 2h at 37°C in the presence of the virus (inoculation condition), for 28h at 37°C after virus removal (post-inoculation condition) or both during the 2h inoculation and 28h post-inoculation periods (inoculation and post-inoculation condition). Cells were fixed with ice-cold methanol and subjected to immunofluorescent detection of viral E1 envelope protein.

### HCV grown in primary culture (HCVpc)

For HCV infection, PHH were inoculated 3 days post-seeding at a MOI of 2 with a non-modified JFH1 virus (HCVcc), as previously described (38). After a 2h incubation at 37°C, the inoculum was removed, and PHH were incubated in complete culture medium for 28h at 37°C at the indicated concentrations of dehydrojuncusol. The medium was then renewed without treatment. Supernatants containing infectious HCVpc were harvested 3 days post-inoculation, and infectious titers were evaluated by a focus-forming assay on Huh-7 cells as previously described (42). Intracellular levels of negative-strand HCV RNA were quantified by a strand-specific reverse-transcription real-time polymerase chain reaction technique described previously (threshold of detection, 25 copies/reaction) (43). The potential cytotoxicity of dehydrojuncusol was assessed by measurement of the activity of lactate dehydrogenase (LDH) released into culture supernatants, as previously described (23).

### BVDV and YFV

Bovine viral diarrhea virus (BVDV, strain NADL) was produced in MDCK cells as previously described (44). Yellow fever virus (YFV, strain 17D) was produced in SW13 cells. Infection assays were performed with MDBK cells (BVDV) or Huh-7 cells (YFV) seeded in 96-well plates, at a MOI of 1.5 or 1, respectively. Cells were infected for 1 hour at 37°C. The viral inoculum was removed and the cells were further cultured for either 15 hours (BVDV) or 23 hours (YFV). Cells were fixed with paraformaldehyde 3% in PBS and subjected to immunofluorescent detection of NS3 (BVDV) or E (YFV).

### HCVpp

Retroviral pseudoparticles expressing HCV envelope glycoproteins E1E2 of genotype 2a (HCVpp) were produced in HEK-293T cells as described (45). Huh-7 cells were seeded in 48-well plates and inoculated with HCVpp in the presence of compounds for 2h. Inoculum was removed and replaced with fresh medium without compound. Cells were lysed at 48h post-inoculation in 50 µl of 1x Luciferase Lysis Buffer (Promega, Madison, USA) and luciferase activity was quantified in a Tristar LB 941 luminometer (Berthold Technologies, Bad Wildbad, Germany) using Luciferase Assay System (Promega) as recommended by the manufacturer.

### Immunofluorescent detection assay

HCVcc, YFV and BVDV infected cells grown in 96-well plates were processed for immunofluorescence detection of viral proteins, as previously described (46). Nuclei were stained with 1 µg/mL of DAPI, and infected cells were detected by immunofluorescent labeling of E1 envelope glycoprotein (HCV), E envelope protein (YFV) and NS3 protein (BVDV) followed by Cy3-conjugated anti-mouse secondary antibody. For BVDV and YFV infection assays, the plates were observed with an Axiophot microscope equipped with a 10×magnification objective (Carl Zeiss AG, Oberkochen, Germany). Fluorescent signals were collected with a Coolsnap ES camera (Photometrix, Kew, Australia). Images of infected cells (Cy3 channel) and of nuclei (DAPI channel) present in 10 randomly picked areas from each well were recorded. The number of total cells and infected cells were quantified using ImageJ software. For quantification of HCVcc infection, confocal images were recorded with an automated confocal microscope IN Cell Analyzer 6000 (GE Healthcare Life Sciences) using a 20× objective with exposure parameters 405/450 nm and 561/610 nm. Six fields per well were recorded. Each image was then processed using the Colombus image analysis software (Perkin Elmer). Nuclei were first segmented and the cytoplasm region was extrapolated based on the DAPI staining. Objects with a specific and predefined size were defined as cells. The ratio of infected cells over total cells represents the infection rate. For each virus, the number of cells per well and MOI were determined in order to have 30 to 40% of infected cells in control experiments with no inhibitor. Infection rates in DMSO controls were expressed as 100%.

### Viral RNA quantification

Huh-7 cells seeded in 12-well plates were inoculated with HCVcc for 2h. The inoculum was removed and replaced with culture medium. Forty-eight hours after inoculation, cells were lysed in lysis buffer from the kit NucleoSpin® RNA Plus (Macherey Nagel), and total RNA extracted following manufacturer’s instructions, eluted in a final volume of 60 µl of H_2_O, and quantified. Ten microliters of RNA were used for cDNA synthesis using High Capacity cDNA Reverse Transcription kit (Applied Biosystems). Five microliters of cDNA were used for real-time reverse-transcription polymerase chain reaction (qRT-PCR) assay using Taqman probes as described previously (47).

### Viability Assay

Huh-7 cells were plated in 96-well plates at a density of 6.000 cells/well and then were incubated the next day in 100 μl of culture medium containing increasing concentrations of dehydrojuncusol for either 24h, 48h, or 72h. An MTS [3-(4,5-dimethylthiazol-2-yl)-5-(3-carboxymethoxyphenyl)-2-(4-sulfophenyl)-2H-tetrazolium]-based viability assay (Cell Titer 96 Aqueous non-radioactive cell proliferation assay, Promega) was performed as recommended by the manufacturer. The absorbance of formazan at 490 nm is detected using an ELISA plate reader (ELX 808 Bio-Tek Instruments Inc). Each measure was performed in triplicate.

### HCV replicon

The plasmid pSGR-JFH1 encoding a sub-genomic replicon of JFH-1 strain was obtained from Dr T Wakita (48). A *Bgl*II and an *Nsi*I restriction sites were inserted between codons Pro419 and Leu420 of NS5A, and the coding sequence of EGFP was then inserted between these two sites. This position was previously shown to accept a GFP insertion in a sub-genomic replicon of the Con1 strain (49). Mutations Leu31Met (L31M) and Tyr93His (Y93H) were inserted independently by PCR. Plasmid pSGR-JFH1-NS5AGFP and PCR amplified DNA containing mutated sequences were digested with *Nsi*I and *Bgl*II restriction enzymes, and PCR fragment inserted by ligation into the plasmid. All constructs were verified by sequencing. These plasmids were *in vitro* transcribed before electroporation into Huh-7 cells. Cells that express wild type and mutated replicons were selected for using 500 µg/mL of geneticin during 15 days and cultured in a medium containing 250 µg/mL of geneticin.

### Replication assay

Huh-7 cells stably expressing wild type or mutated replicons were seeded in 24-well plates and incubated with the different compounds for 24h, 48h and 72h. They were lysed in ice cold lysis buffer (Tris HCl 50mM, NaCl 100 mM EDTA 2 mM Triton-100 1% SDS 0.1%) on ice for 20 min. Lysates were collected and 20 µg of proteins were analyzed by western blotting using anti-NS5A and anti-β tubulin antibodies. Peroxidase-conjugated goat anti-mouse secondary antibody (Jackson Immunoresearch) was used for the revelation using ECL western blotting substrate (Thermo Fischer Scientific). The intensity of the bands was quantified using ImageJ software.

Wild-type, L31M and Y93H replicon cell lines were seeded in 96-well plates and incubated with boceprevir, daclatasvir or methylene chloride partition at given concentration for 72h. Cells were fixed with PFA 3% for 30 min. The cells were counted by labeling of nuclei with DAPI (0.5 µg/mL). Number of GFP-positive cells was quantified with a Zeiss Axiophot 2 microscope equipped with a 40×/1.3 numerical aperture lens. Fluorescence signals were analyzed as described previously.

### Selection of a dehydrojuncusol-resistant virus and identification of resistance mutations

Huh-7 cells were infected with JFH1 followed by treatment with 5 × EC_50_ (6.55 µM) of dehydrojuncusol. Cells were split every 3-4 days in the presence of dehydrojuncusol at 6.55 µM and viral RNA was extracted in parallel to follow appearance of resistant viruses. At day 21, viral RNA was extracted and the genomic sequence of HCV from NS2 to NS5B was amplified by RT-PCR and sequenced. Amino acid changes that arose during inhibitor selection were identified by analysis of the DNA sequence compared to the initial and control passages in the absence of drug. The identified mutations were reintroduced into the JFH1 plasmid by PCR mutagenesis, and the plasmids were sequenced.

### Statistical Analysis

The results were presented as means ± SEM of three independent experiments performed in triplicate. The statistical test used is a Kruskal Wallis nonparametric test followed by a Dunn’s multicomparison post hoc test with a confidence interval of 95% to identify individual difference between treatments. P values < 0.05 were considered as significantly different from the control. The data were analyzed using Graph Pad Prism (Version 5.0b) by comparisons between each treated group and untreated group (DMSO control).

## ACKNOWLEDGMENTS

This work was supported by the French National Agency for Research on AIDS and Viral Hepatitis (ANRS-18208), the European Community (ERC-STG INTRACELLTB Grant n° 260901), the Agence Nationale de la Recherche (ANR-10-EQPX-04-01), the Feder (12001407 (D-AL) Equipex Imaginex BioMed) and the Région Nord Pas de Calais (convention n°12000080). The authors wish to thank platforms of CUMA (University of Lille 2, Professor J. F. Goossens) and LARMN (University of Lille 2, Professor N. Azaroual) for access to equipment. We are grateful to J. Bukh, C.M. Rice and T. Wakita for providing essential reagents. The authors are also grateful to Dr. Abderrazak Smaoui (Biotechnology Centre of Borj-Cédria) concerning the botanical identification and to Thibaut Vausselin for useful discussions. M.-E.S. is a recipient of a PhD Fellowship provided by the French Government.

